# COVID-19 and Cholinergic Anti-inflammatory Pathway: *In silico* Identification of an Interaction between α7 Nicotinic Acetylcholine Receptor and the Cryptic Epitopes of SARS-CoV and SARS-CoV-2 Spike Glycoproteins

**DOI:** 10.1101/2020.08.20.259747

**Authors:** George Lagoumintzis, Christos T. Chasapis, Nikolaos Alexandris, Socrates Tzartos, Elias Eliopoulos, Konstantinos Farsalinos, Konstantinos Poulas

## Abstract

SARS-CoV-2 is the coronavirus that originated in Wuhan in December 2019 and has spread globally. The observation of a low prevalence of smokers among hospitalized COVID-19 patients has led to the development of a hypothesis that nicotine could have protective effects by enhancing the cholinergic anti-inflammatory pathway. Based on clinical data and on modelling and docking experiments we have previously presented the potential interaction between SARS-CoV-2 Spike glycoprotein and nicotinic acetylcholine receptors (nAChRs), due to a “toxin-like” epitope on the Spike Glycoprotein, with homology to a sequence of a snake venom toxin. We here present that this epitope coincides with the well-described cryptic epitope for the human antibody CR3022 and with the epitope for the recently described COVA1-16 antibody. Both antibodies are recognizing neighboring epitopes, are not interfering with the ACE2 protein and are not able to inhibit SARS-CoV and SARS-CoV-2 infections. In this study we present the molecular complexes of both SARS-CoV and SARS-CoV-2 Spike Glycoproteins, at their open or closed conformations, with the molecular model of the human α7 nAChR. We found that the interface of all studied protein complexes involves a large part of the “toxin-like” sequences of SARS-CoV and SARS-CoV-2 Spike glycoproteins and toxin binding site of human α7 nAChR.

## 1. INTRODUCTION

Severe acute respiratory syndrome (SARS) is a viral respiratory disease caused by the SARS-associated coronavirus (SARS-CoV). In February 2003, SARS was first reported in Asia. The disease spread to countries in North America, South America, Europe, and Asia before the SARS global outbreak of 2003 was contained. No known cases of SARS were reported anywhere in the world after 2004 [1,2]. In April 2003, rumors spread that smoking protected patients from developing SARS, which were rejected by the authorities [3]. Several studies suggested that smokers were under-represented among hospitalized SARS patients but no firm conclusions about the association between smoking and SARS-CoV infection were drawn [4-7].

As the global pandemic of Corona Virus Disease 2019 (COVID-19), a disease caused by the acute respiratory syndrome coronavirus 2 (SARS-CoV-2), is evolving, it is imperative to understand the pathophysiology and the risk and potentially protective factors associated with disease progression and severity in order to offer effective therapies. The association between smoking and COVID-19 is controversial. Several meta-analyses have identified an unusually low pooled prevalence of smoking compared to the population smoking rates [8, 9]. While limitations exist in these analyses, similar observations of low smoking prevalence among hospitalized COVID-19 patients have been observed in case series from many countries [10]. Cox et al. recently published a cohort study of 8.28 million participants, including 19,486 confirmed COVID-19 cases, and found that smoking was associated with lower odds for COVID-19 diagnosis and ICU admission [11]. This apparently protective effect was stronger for heavy and moderate smokers. Other studies have shown that, once hospitalized, smokers are at higher risk for adverse outcomes [9,12]. We hypothesized for the first time in early April 2020 that the Nicotinic Cholinergic System (NCS) could be involved in the pathophysiology of severe COVID-19 and we have recently expanded on this hypothesis [13,14].

The NCS is an important pathway which regulates the response to inflammation. Its effects on macrophages and other immune cells are mainly regulated by the vagus nerve and by alpha7 nicotinic acetylcholine receptors (nAChRs) [15]. This so-called “cholinergic anti-inflammatory pathway” has been found beneficial in preventing inflammatory conditions such as sepsis and Acute Respiratory Distress Syndrome (ARDS) in animal models [16]. Dysregulation of the NCS could therefore be a possible cause for the uncontrolled inflammatory response in COVID-19. It could also explain other clinical manifestations of COVID-19 such as anosmia and thromboembolic complications [17]. We therefore hypothesized that SARS-CoV-2 may interact directly with the NCS [18]. Through computational modeling and docking experiments, we have identified a key interaction between the SARS-CoV-2 Spike glycoprotein, mainly amino acids (aa) 381-386, and the nAChR alpha subunit (mainly aa 189-192) extracellular domain (ECD), a region that forms the core of the nAChRs “toxin-binding site”. Similar to the interaction between the alpha subunit of nAChR and α-bungarotoxin, this interaction reinforced the possibility of SARS-CoV-2 interacting with nAChRs [18]. It is interesting that other research groups have identified such an interaction through different epitopes of SARS-CoV-2 Spike glycoprotein [19].

Yuan et al. recently presented the crystal structure of CR3022, a human antibody previously isolated from a convalescent SARS patient, in complex with the receptor binding domain (RBD) of the SARS-CoV-2 Spike (S) glycoprotein at 3.1Å resolution [20]. Huo et al. also reported that CR3022 binds to the RBD, and presented the crystal structure of the Fab/RBD complex at 2.4Å resolution [21]. The antibody’s neutralizing potential is established for SARS-CoV but not for SARS-CoV-2, although it may exhibit some *in vivo* protection against the latter. Both groups recognized that the highly conserved and cryptic epitope for CR3022 is inaccessible in the prefusion Spike glycoprotein. Neither group could explain the exact function of this epitope, but both proposed that this cryptic epitope could be therapeutically useful, possibly in synergy with an antibody that blocks ACE2 attachment. The aa that form this epitope are highly conserved in SARS-CoV and SARS-CoV-2. Of the 28 epitope residues (defined as residues buried by CR3022), 24 are preserved between SARS-CoV and SARS-CoV-2. This explains the CR3022 cross-reactivity to SARS-CoV and SARS-CoV-2 RBDs. Nevertheless, despite having high epitope residue conservation, CR3022 Fab binds to SARS-CoV RBD with much greater affinity than to SARS-CoV-2 RBD. CR3022’s epitope does not overlap with the ACE2-binding site. Structural alignment of the CR3022-SARS-CoV-2 RBD complex with the ACE2-SARS-CoV-2 RBD complex further implies that CR3022 binding does not interfere with ACE2 [20]. This suggests that the neutralization mechanism of CR3022 for SARS-CoV is not based on direct blocking of receptor binding, which is consistent with the finding that CR3022 does not compete with ACE2 for binding to the RBD [22]. Liu et al. recently presented a SARS-CoV-2 specific antibody, called COVA1-16, isolated from a convalescent COVID-19 patient, which binds to the same epitope [23]. The affinity for SARS-CoV-2 Spike glycoprotein is higher, compared to the affinity for SARS-CoV Spike glycoprotein, as expected.

In this study, we examined the potential interaction between α7 nAChRs and SARS-CoV and SARS-CoV-2 Spike glycoproteins RBDs. For SARS-CoV-2 our previous study identified such an interaction located at aa 375-390, a region that is homologous to a sequence of the NL1 homologous neurotoxin which is a well-established NCS inhibitor [24,25]. Interestingly this epitope is part of the cryptic epitope mentioned above for the human antibody CR3022 and COVA1-16 [20,21,23]. Having previously presented the 3D structural location of this “toxin-like” sequence on the Spike glycoprotein and the superposition of the modelled structure of the Neurotoxin homolog NL1 and the SARS-CoV-2 Spike glycoprotein [18], we are extending our previously published molecular modeling and docking experiments by presenting the complexes of both SARS-CoV and SARS-CoV-2 S glycoproteins with the ECD of the model of human α7 nAChR pentamer, in their “open” and “closed” conformations, ideally adopted by Spike glycoproteins.

## 2. METHODS

### 2.1. Sequence retrieval and alignment

We compared aa sequences between SARS-CoV and SARS-CoV-2 Spike glycoproteins and snake venom neurotoxins. Sequence retrieval of the protein sequences of both virus-related Spike proteins and “three finger” neurotoxins from various species was performed using the National Center for Biotechnology Information (NCBI, Bethesda, MD, USA) databases. BLAST searches were performed using Mega BLAST [26] with the UNIPROT protein database [27] and by using BLASTP (protein–protein BLAST) with default parameters. Multiple sequence alignments were performed using ClustalOmega program (Clustal-O) [28].

### 2.2. Structure retrieval

The 3D structures of the SARS-CoV-2 Spike glycoprotein (S1) in complex with the human Angiotensin Converting Enzyme 2 (hACE2) (PDB id: 6LZG), the hACE2 (1R41, 1R42) the cryo-EM determined complex of spike protein S-ACE2-B0AT1 neutral amino acid transporter (PDB id: 6M18), the structure of a neutralizing to SARS-CoV mAb that also cross-reacts with the S protein of SARS-CoV-2 when the latter is in complex with the ACE2 receptor (PDB id: 6NB7), the ECD of the nAChR α9 subunit in complex with α-bungarotoxin (PDB id: 4UY2), the structure of SARS-CoV-2 RBD in complex with the human monoclonal antibody CR3022 (PDB id:6W41) and the structure of the ligand binding domain (LBD) of a chimera pentameric α7 nAChR (PDB id: 3SQ9) were downloaded from the Protein Data Bank (PDB).

### 2.3. Molecular modelling and docking experiments

The protein structure prediction of the ECD of human α7 nAChR was performed using ROSETA software [29], applying automated multi-step and multi-template homology modelling approach. The complexes between SARS-CoV and SARS-CoV-2 Spike 1 (S1) glycoprotein and ECD of the human pentameric α7 nAChR were modelled using HADDOCK server [30]. The predicted interaction surfaces and the ambiguous interaction restrains (AIRs) which were used to drive HADDOCK process were automatically generated using WHISCY software. The input data for WHISCY prediction of the interaction surface between ECD of human pentameric α7 nAChR and SARS-CoV and SARS-CoV-2 S1 glycoproteins were the conserved residues that were experimentally identified to be involved in the interaction between α7 nAChR chimera and α-bungarotoxin. The binding affinity of biomolecular complexes were predicted using PRODIGY software [31]. All the protein structures are visualized using UCSF chimera software [32].

## 3. RESULTS

### 3.1. Sequence alignment

**Figure 1A** presents the sequence alignment of SARS-CoV and SARS-CoV-2 S1 glycoproteins (A7J8L4, P0DTC2) with Neurotoxin homolog NL1 (Q9DEQ3). We found a double “recombination” within the same S-protein sequence (aa 375-390), which is homologous in the Neurotoxin homolog NL1 sequence, part of the “three-finger” interacting motif of the toxins. This peptide fragment (aa 375-390) is part of the RBD (aa 319-541) of the SARS-CoV and SARS-CoV-2 Receptor Binding Domain (RBD) (aa 306-527) Spike glycoproteins (the domain through which the spike protein recognizes the ACE2 on the host’s cell surface) neighboring to the ACE2 Receptor Binding Motif (aa 437-508). Quite importantly, this peptide is the main part of the epitope for the CR3022 antibody, as described before [20,21]. The main interacting amino acids between the RBD of SARS-CoV and mAb CR3022, as described in the crystal structure of CR3022 and SARS-CoV-2 Spike glycoprotein [20], are shown in Figure 1B. Molecular models of mAb CR3022 interacting with SARS-CoV, are presented in Figure 1C.

**Figure 1.**
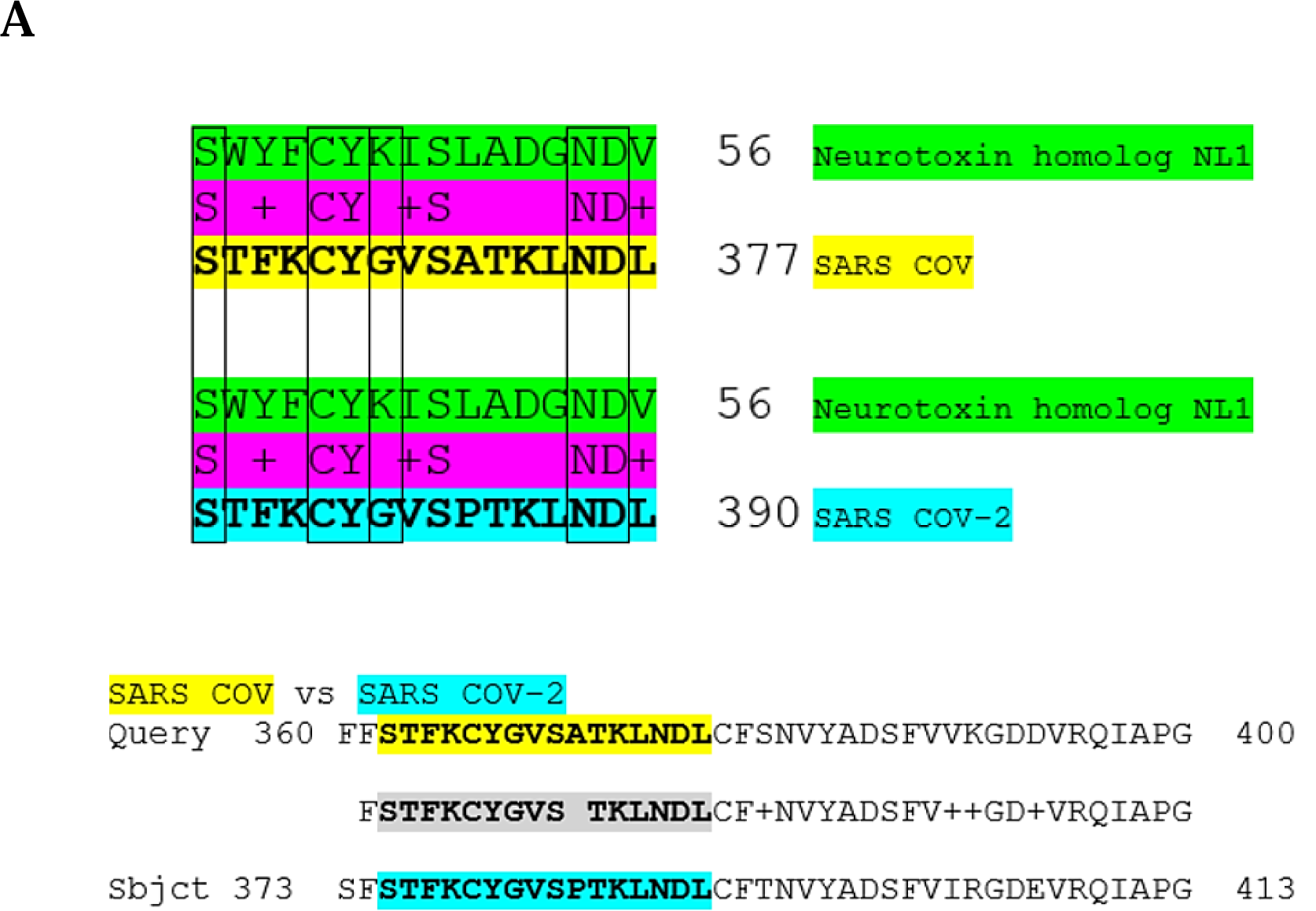
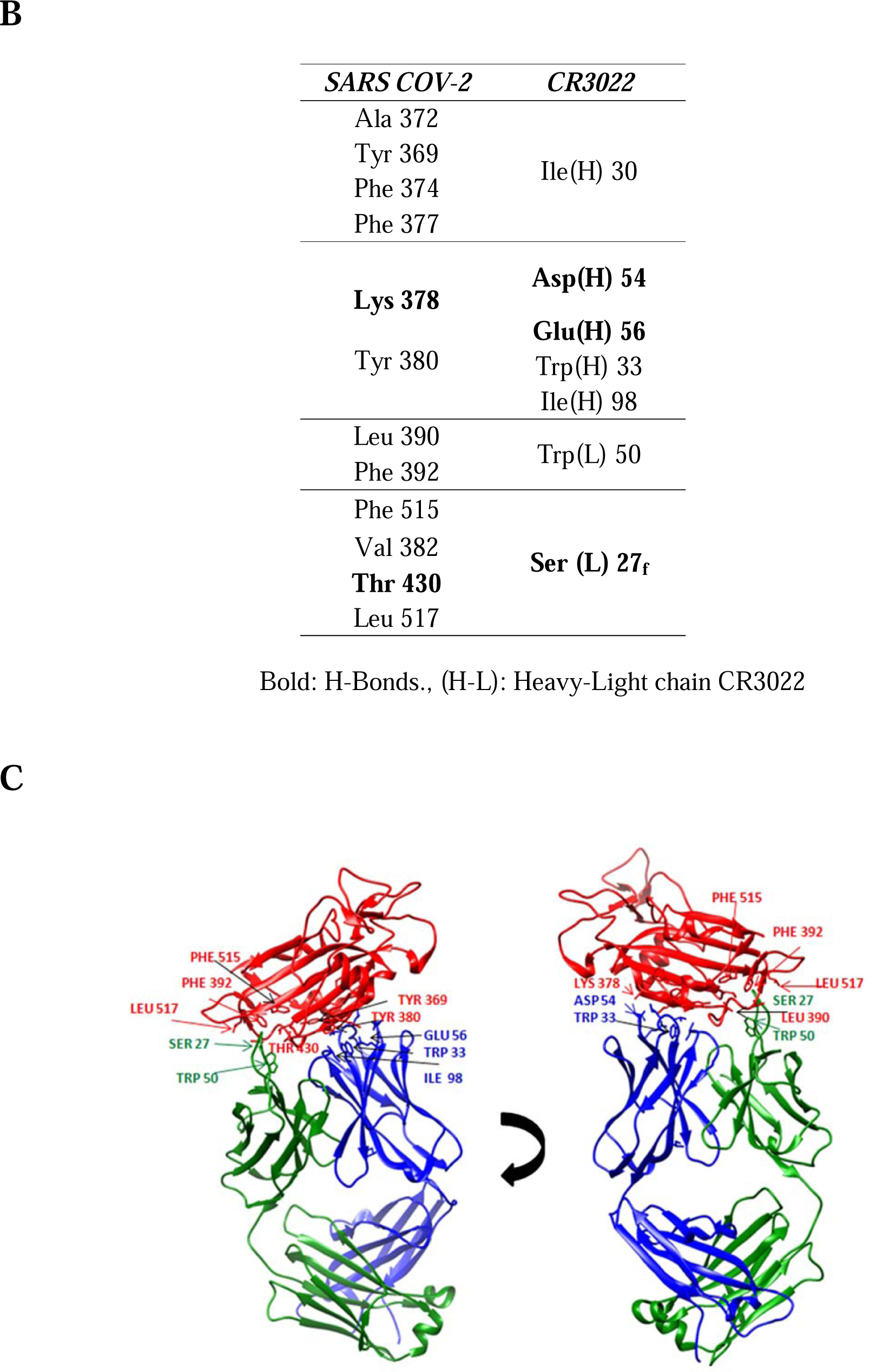
**(A)** Sequence alignment between the SARS-CoV and SARS-CoV-2 Spike glycoproteins and Neurotoxin homolog NL1 depicting amino acids within this sequence which are identical or functionally equivalent to Neurotoxin homolog NL1 toxin. **(B)** Amino acid interactions between SARS-CoV-2 RBD and SARS-CoV neutralizing mAb CR3022. **(C)** Spatial amino acid interactions between SARS-CoV-2 RBD (red) and CR3022 heavy (blue) and light chain (green) through different angles.

### 3.2. Molecular modelling of toxin-like fragment of SARS-CoV and SARS-CoV-2 RBD

This “toxin-like” fragment on SARS-CoV (aa 362-377) and SARS-CoV-2 (aa 375-390) RBD, containing an amphipathic sequence of alternating polar and hydrophobic amino acid residues with selectively charged amino acids in a conserved order, lies on the spike protein surface and is not buried in the domain core. The toxin-like sequence, in ball and stick representation, and its location in the protein surface is illustrated in **Figure 2**. Neighboring the ACE2 binding motif, this entity may interact with the human α7 nAChRs in a manner similar to neurotoxins.

**Figure 2.**
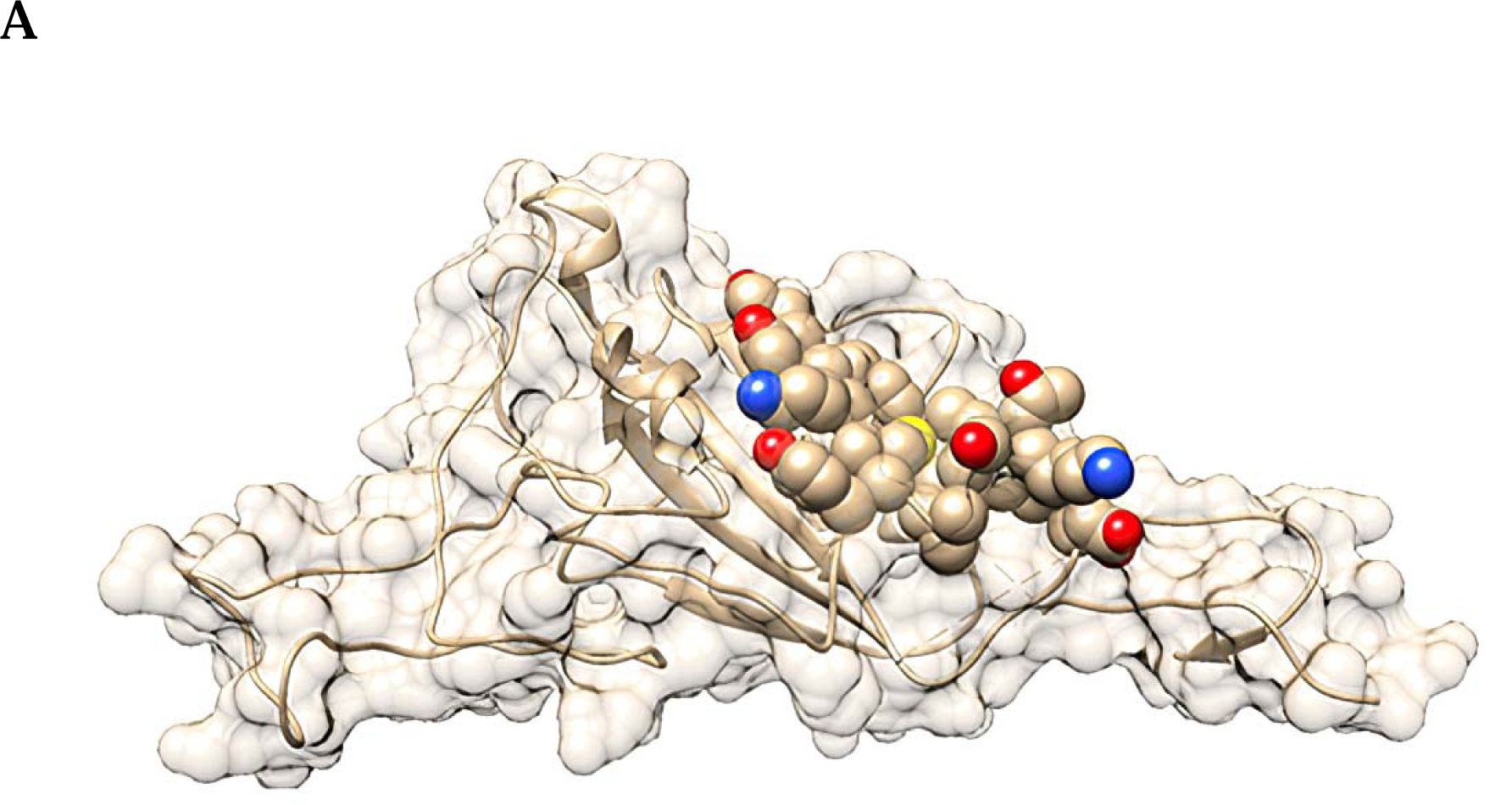
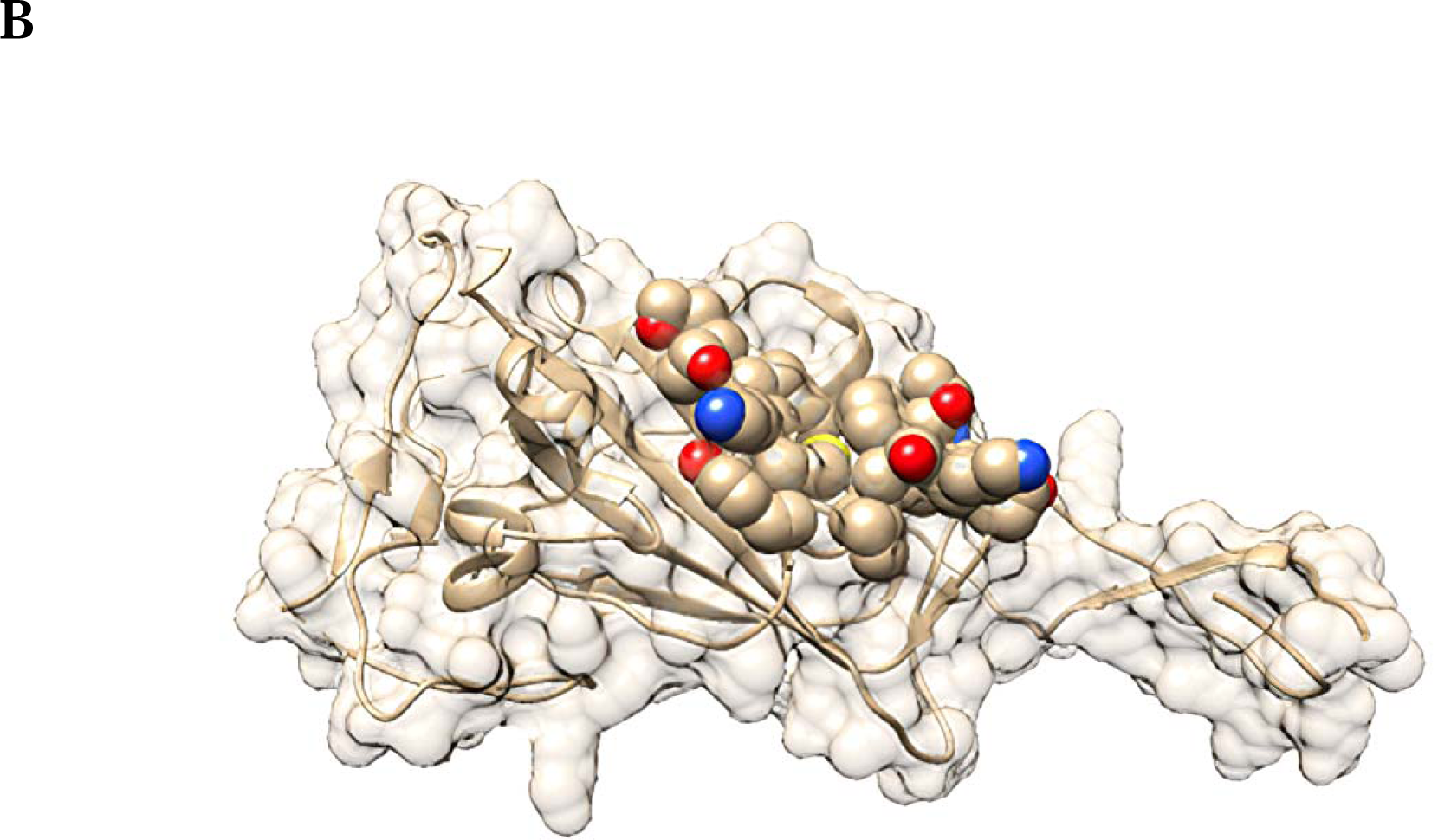
Structural location of the toxin-like sequence within the SARS-CoV **(A)** and SARS-CoV-2 **(B)** Spike glycoprotein. The toxin-like sequence is illustrated in ball and stick format.

### 3.3. Interaction of SARS-CoV and SARS-CoV-2 S1 with the ECD of human α7 nAChR

We have previously identified the interaction between the SARS-CoV-2 S1 glycoprotein (aa 381-386) and the α9 subunit of nAChR ECD (aa 189-192), a region that forms the core of the nAChR “toxin-binding site”. The interaction between the two proteins is caused by complementarity of the hydrogen bonds and shape [18]. The interaction mode is very similar to the interaction between α9 nAChR and both α-bungarotoxin and the homologous neurotoxin NL1 (snake venom toxins are known to inhibit nAChRs). Similar interacting surfaces were observed between the SARS-CoV and SARS-CoV-2 S1 and the LBD of the pentameric α7 nAChR chimera.

Herein, the HADDOCK models show that the interface of all studied protein complexes involves the majority of the toxin like sequences within SARS-CoV S proteins and toxin binding site of human α7 nAChR. The binding affinity (ΔG, expressed in kcal mol^-1^), the dissociation constant (K_d_ at 25 □, expressed in Molar), electrostatic energy (expressed in kcal mol^-1^) and the buried surface area (expressed inÅ^2^) for all the modelled protein complexes are presented in **Table 1**. The dissociation constant of all SARS-α7 nAChR complexes found to be in the nM range, which is comparable with experimental supported Kds of well-known enzymatic interacting partners that produce stable protein complexes (*i*.*e*., E2-E3 pairs in ubiquitination pathway [33]).

**Table 1.** Haddock parameters of SARS-CoV S1 and SARS-CoV-2 S1 with ECD of human α7 nAChR pentamer.

|  | SARS-CoV S1 |  | SARS-CoV-2 S1 |  |
| --- | --- | --- | --- | --- |
|  | *Open* | *Closed* | *Open* | *Closed* |
| | with human α7 nAChR | | | |
| **$\Delta G$ (kcal mol<sup>-1</sup>)** | -8,5 | -10,6 | -10 | -9,4 |
| **Kd (Molar) at 25.0 °C** | 5.6E-07 | 1.6E-08 | 4.6E-08 | 1.3E-07 |
| **HADDOCK score** | -117.6 (8.5) | -96.2 (4.4) | -38.9 (12.0) | -46.2 (7.9) |
| **Electrostatic Energy (kcal mol<sup>-1</sup>)** | -174.4 (31.4) | -114.6 (28.9) | -198.9 (19.6) | -41.6 (14.5) |
| **Buried Surface Area (Å<sup>2</sup>)** | 1777.7 (251.6) | 1845.9 (109.3) | 1561.8 (60.3) | 1677.1 (122.5) |

Figure 3 shows the clusters of intermolecular contacts (ICs) at the interface (within the threshold distance of 5.5 □) for the complexes between SARS-CoV S1 **(A)** and SARS-CoV-2 S1 **(B)** glycoproteins and the LBD of the human α7 nAChR. The main ICs cluster for each interaction, involves the regions: ^355^Phe-^372^Cys, ^365^Cys-^375^Ser and ^383^Ser-^388^Cys and ^207^Glu- ^217^Tyr of SARS-CoV, SARS-CoV-2 S1 protein and the LBD of the human α7 nAChR, respectively.

The HADDOCK models show that the interface of all studied protein complexes involve the majority of the toxin like sequences within SARS-CoV Spike glycoproteins and toxin binding site of human α7 nAChR. The relative orientation of RBD Spike glycoprotein in SARS-CoV and LBD of α7 nAChR differs significant between the open and closed conformation of their complexes (**Figure 4**).

**Figure 3.**
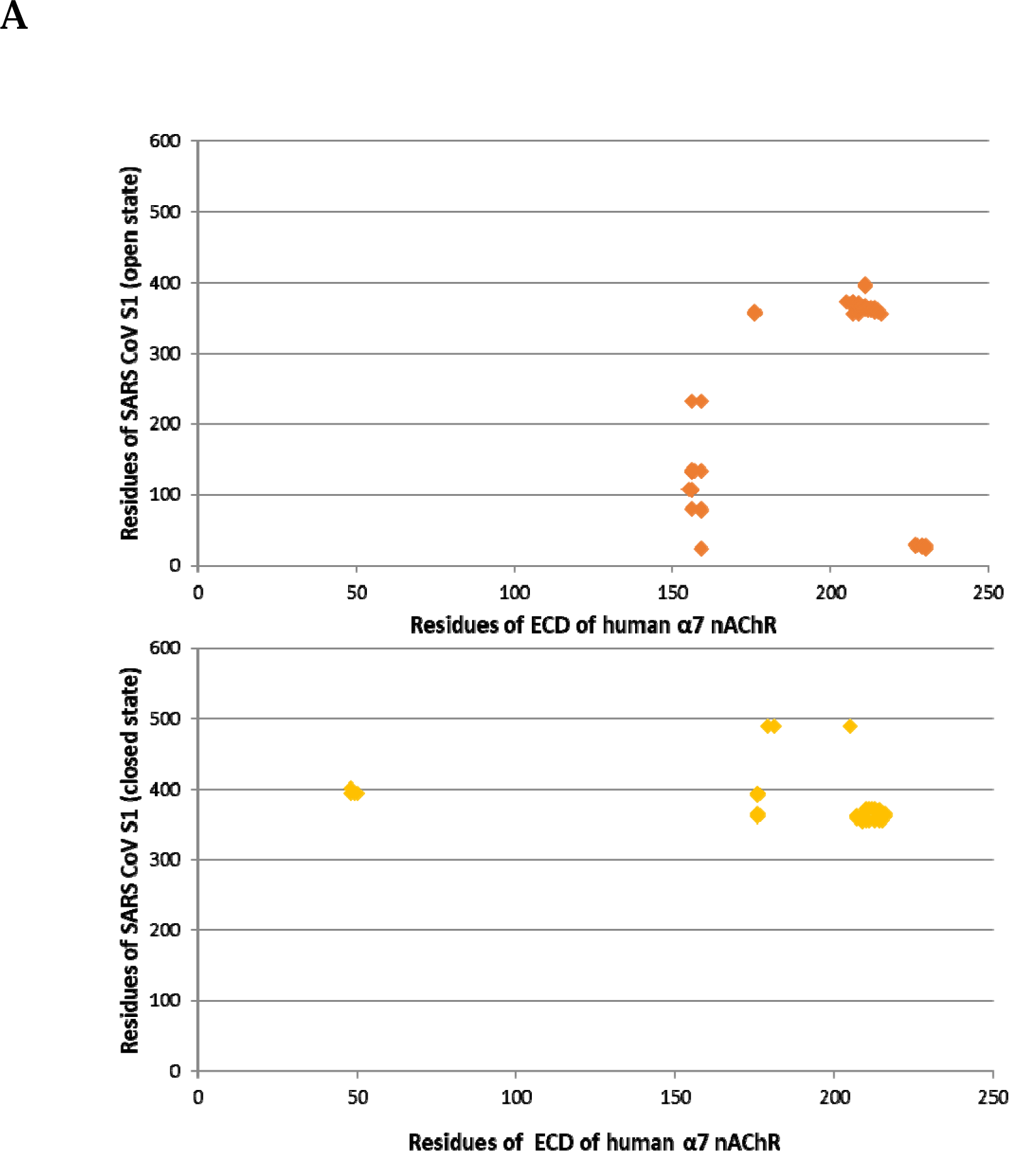
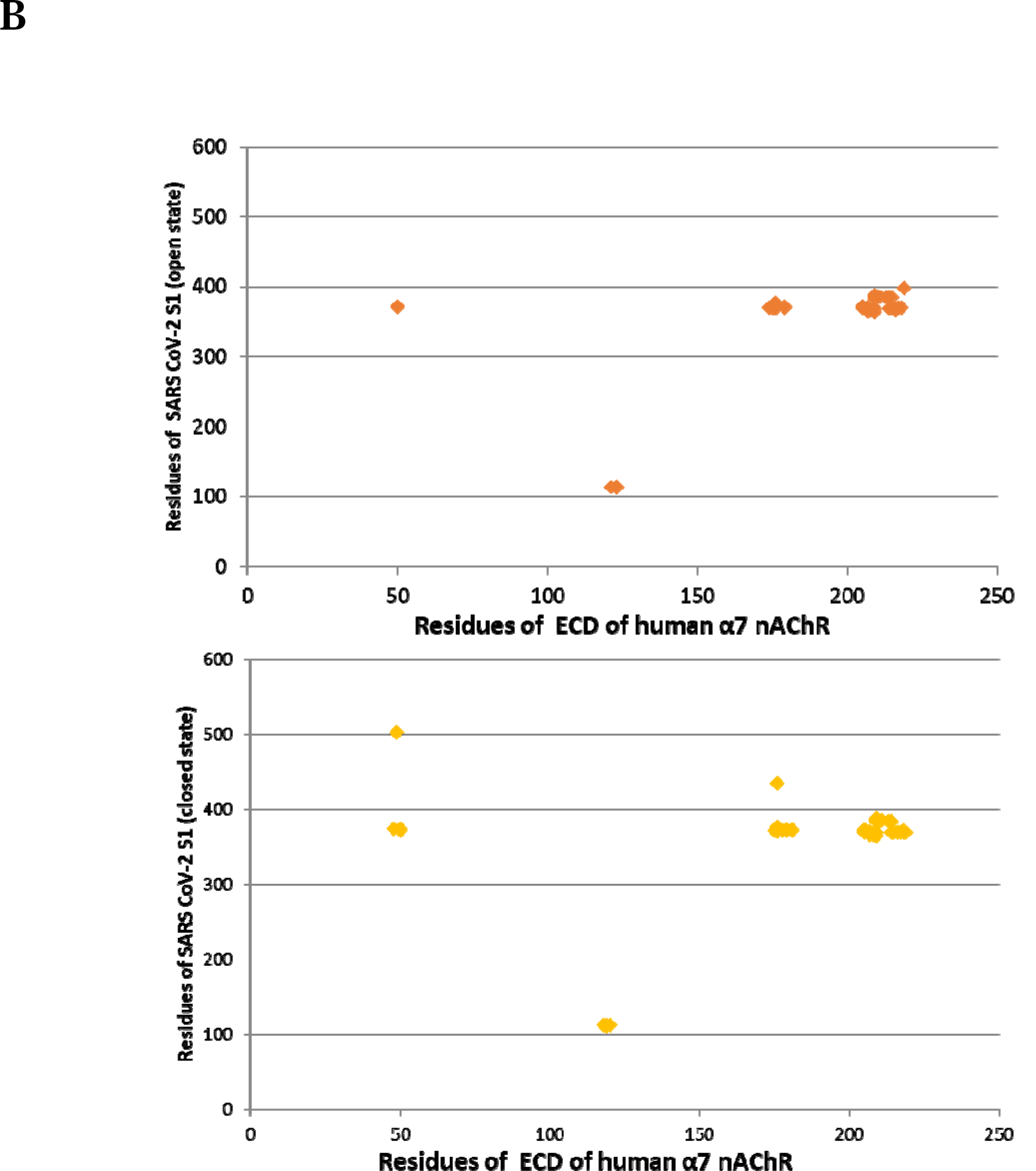
Cluster of intermolecular contacts (ICs) at the interface (within the threshold distance of 5.5 □) for the complexes between SARS-CoV S1 **(A)** and SARS-CoV-2 S1 **(B)** glycoproteins in open and closed conformation with the LBD of the human ECD of α7 nAChR.

**Figure 4.**
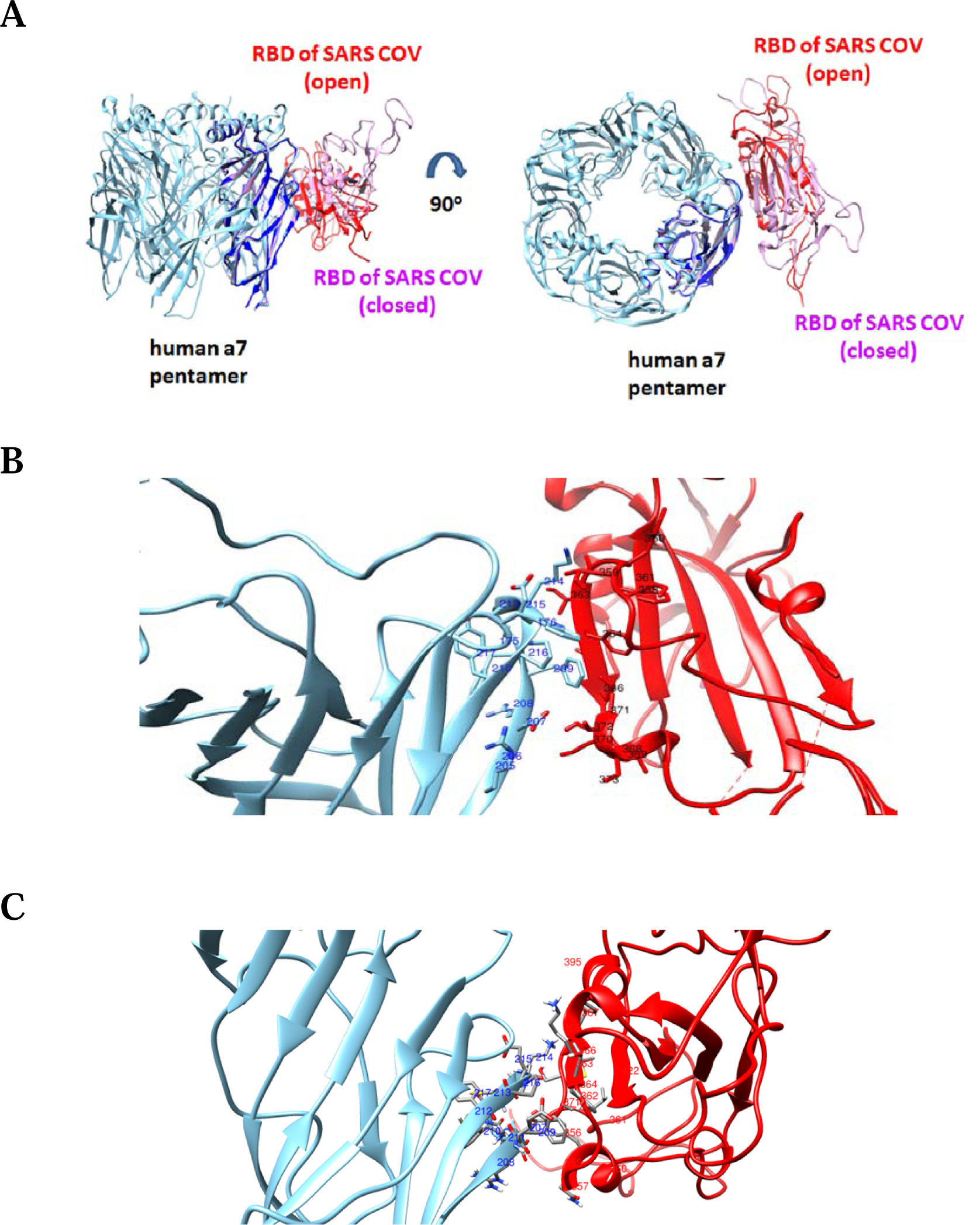
**(A)** HADDOCK complexes of the RBD of SARS-CoV in its open and closed state with one subunit of the pentameric extracellular domain of human α7 nAChR. (**B)** The interaction interface between the RBD of SARS-CoV in open state (red color) with the extracellular domain of human α7 nAChR (cyan color). (**C)** The interaction interface between the RBD of SARS-CoV in closed state (red color) with the extracellular domain of human α7 nAChR (cyan color).

Slightly different orientations of the protein interactions were observed between the open and closed complexes of SARS-CoV-2 Spike with the LBD of α7 nicotinic receptor (**Figure 5**). In both open and closed conformation, the orientation of the RBD of SARS CoV-2 when interacting with the α7 subunit is more similar with the interaction of the closed, rather than the open, conformation of the RBD of SARS CoV.

**Figure 5.**
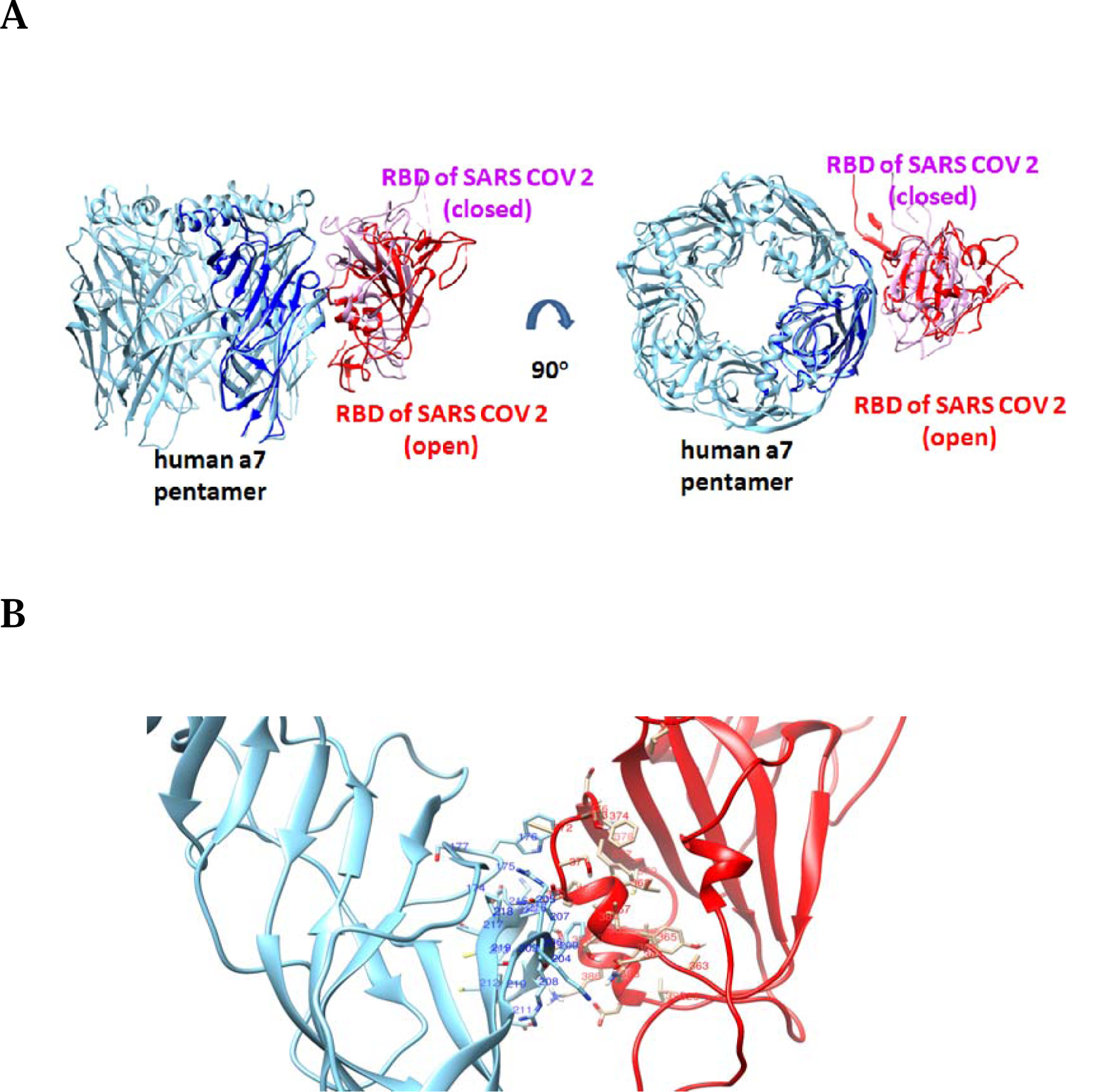
(**A)** HADDOCK complexes of RBDs of SARS-CoV-2 in their open and closed state with one subunit of the pentameric extracellular domain of human α7 nAChR. (**B)** The interaction interface between RBD Spike of SARS-CoV-2 in open state (red color) with extracellular domain of human α7 nAChR (cyan color).

Detailed interacting surfaces of SARS-CoV and SARS-CoV-2 S1 RBDs and α7 nAChRs complexes are illustrated in the Figures 4B, 4C and 5B. Two residues which are conserved in SARS cryptic epitopes (^366^Cys and ^367^Tyr) are critical for their interaction with the α7 subunit (Figure 6). Specifically, SARS CoV-α7 contacts ^366^Cys - ^214^ Lys and ^367^Tyr - ^214^Lys in closed conformation, and ^366^Cys -^209^Phe and ^367^Tyr - ^211^Glu in open conformation were identified (**Figure 6**). Also, the ^384^Pro and ^385^Thr, which are conserved residues in the SARS CoV-2 cryptic epitope, are in closed contact with ^214^Lys and ^209^ Phe of the α7 subunit (**Figure 7**).

**Figure 6.**
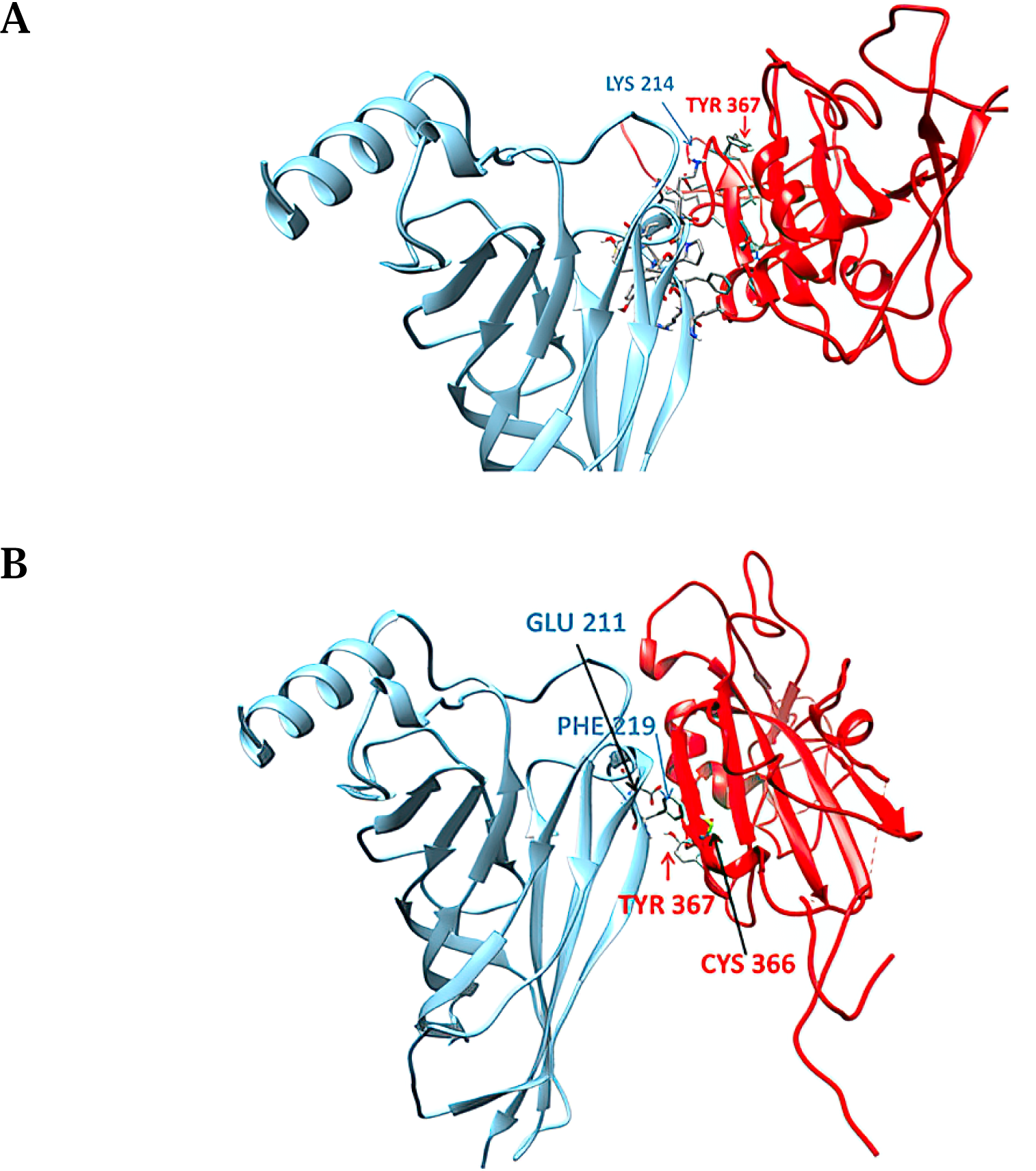
Contacts between ^366^Cys and ^367^Tyr of SARS-CoV cryptic epitope with residues of the α7 subunit in closed **(A)** and open **(B)** state of their complexes.

**Figure 7.**
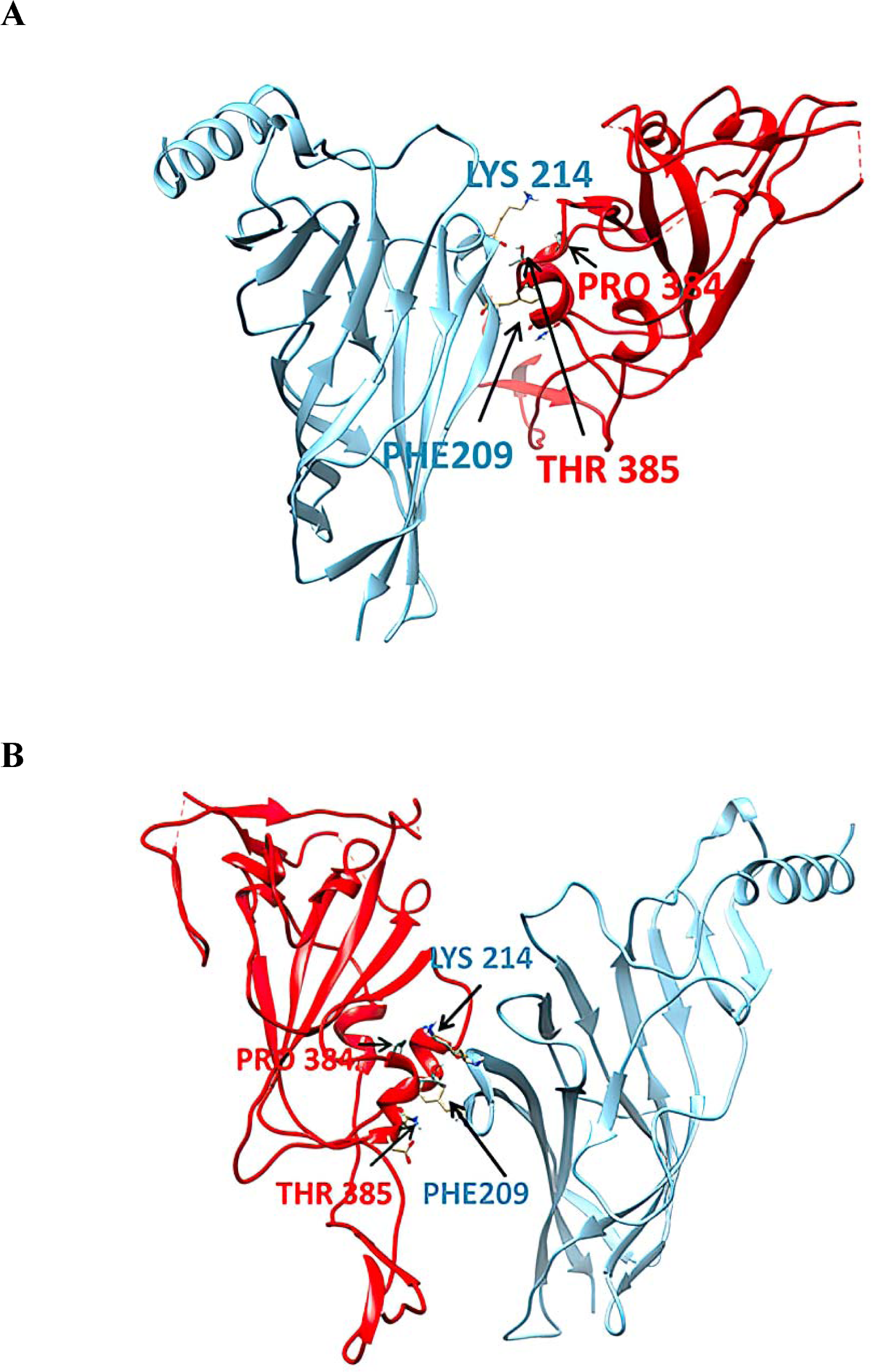
Contacts between ^384^Pro and ^385^Thr of SARS-CoV-2 cryptic epitope with residues of the α7 subunit in closed **(A)** and open **(B)** state of their complexes.

## 4. DISCUSSION

According to our hypothesis, SARS-CoV-2 Spike glycoprotein has a “toxin-like” sequence in its RBD which could bind to the toxin-binding domain of the alpha subunit of the nAChRs. This binding could produce a number of adverse effects by dysregulating the NCS, in which α7 nAChRs are mainly involved. One of the effects could be the dysfunction of the cholinergic anti-inflammatory pathway, resulting in cytokine storm and failure for the immune response to return to homeostasis. Several clinical manifestations of COVID-19 could be explained by cholinergic dysfunction [34]. These findings, if proven *in vivo*, could have important therapeutic implications.

Herein we extent our previous findings of an interaction between SARS-CoV-2 Spike glycoprotein (aa 381-386) and the nAChR α9 subunit, by presenting the complexes of both SARS-CoV and SARS-CoV-2 Spike glycoproteins with the ECD of the model of the human α7 nAChR pentamer, in their “open” and “closed” conformations.

Recently, Yan et al. characterized a conserved cryptic epitope on the Spike glycoprotein, recognized by the human mAb CR3022, which does not appear to inhibit the binding to the ACE2 protein [20]. Similarly, Liu et al. identified and characterized the COVA1-16 mAb, which binds to SARS-CoV-2 Spike glycoprotein to a similar region [23]. Most RNA viruses through antigenic mutation can avoid adaptive immunity of the host [35]. This is achieved due to the error-prone nature of RNA polymerases which induces sequence variation in each virus cycle replication, allowing frequent generation of mutant species [36]. Therefore, naturally occurring “epitopes” in viral antigens, usually targeted by neutralizing antibodies, are intrinsically subjected to antigenic variations. However, some proteins sites on the viral surface are less susceptible to mutation [37-39]. Such conserved regions are typically related to functions that are necessary for viral infection, such as receptor binding or membrane fusion, and are, therefore, less affected by antigenic variability [40].

Quite surprisingly, the cryptic epitope on the SARS-CoV and SARS-CoV-2 Spike glycoproteins is highly conserved and coincides with the toxin-like sequence interacting with nAChR. According to this hypothesis, CR3022, mutants of CR3022 with higher affinity to SARS-CoV-2 Spike glycoprotein and the COVA1-16 antibody can inhibit the interaction between Spike Glycoproteins and nicotinic cholinergic system.

## 5. CONCLUSIONS

These *in silico* findings support our recently published hypothesis that SARS-CoV-2 could interact with nAChRs causing dysregulation of the NCS and the cholinergic anti-inflammatory pathway. Such interactions may lead to uncontrolled immune response and cytokine storm, which is implicated in severe COVID-19 pathophysiology. Our efforts towards elucidating the potential molecular interactions between SARS-CoV-2 S glycoprotein and human nAChRs could provide a basis for accelerated vaccine, immunotherapy and diagnostic strategies against SARS-CoV-2 and other associated betacoronaviruses that could pose a major global human threat in the future.

## ACKNOWLEDGEMENTS

Authors are grateful to the “National Research Infrastructures on Integrated Structural Biology, Drug Screening Efforts and Drug Target Functional Characterization (INSPIRED)” for financial support of the personnel.

## AUTHOR CONTRIBUTIONS

**GL:** *Methodology, Investigation, Software, Writing-Reviewing and Editing;* **CC:** *Methodology, Investigation, Software, Writing-Reviewing and Editing;* **NA:** *Methodology, Investigation, Software, Writing-Reviewing and Editing;* **ST**: *Methodology, Writing-Reviewing and Editing* **EE:** *Methodology, Software, Supervision;* **KF:** *Conceptualization, Reviewing results and data interpretation, Writing Original and final draft;* **KP:** *Writing-Reviewing and Editing, Conceptualization, Supervision*.

**GL, CC**, *and* **NA** contributed equally to this work.

## Abbreviations

aa: Amino Acids
ARDS: Acute Respiratory Distress Syndrome
ACE2: Angiotensin Converting Enzyme 2
COVID-19: Corona Virus Disease 2019
ECD: Extracellular Domain
LBD: Ligand-binding domain
nAChR: Nicotinic Acetylcholine Receptor
NCS: Nicotinic Cholinergic System
RBD: Receptor Binding Domain
SARS-CoV: Severe Acute Respiratory Syndrome Coronavirus
SARS-CoV-2: Severe Acute Respiratory Syndrome Coronavirus 2.

## REFERENCES

1. Center for Disease Control and Prevention, https://www.cdc.gov/sars/index.html

2. Umesh D Parashar, Larry J Anderson, Severe acute respiratory syndrome: review and lessons of the 2003 outbreak, International Journal of Epidemiology, Volume 33, Issue 4, August 2004, Pages 628–634, https://doi.org/10.1093/ije/dyh198.

3. CNN. Hong Kong squashes SARS smoking ‘cure’. [cited 2003 Apr 20]. Available at: https://edition.cnn.com/2003/WORLD/asiapcf/east/04/18/china.sars.smoking/ (accessed on August 14, 2020).

4. Rainer, T., Smit, D., & Cameron, P. (2004). Smoking and Severe Acute Respiratory Syndrome. Hong Kong Journal of Emergency Medicine, 11(3), 143–145. doi. 10.1177/102490790401100303.

5. Tsui PT, Kwok ML, Yuen H, Lai ST. Severe acute respiratory syndrome: clinical outcome and prognostic correlates. Emerg Infect Dis. 2003;9(9):1064–1069. doi: 10.3201/eid0909.030362.

6. Tsang K, Ho P, Ooi G et al. A Cluster of Cases of Severe Acute Respiratory Syndrome in Hong Kong N Engl J Med 2003; 348:1977–1985 doi: 10.1056/NEJMoa030666.

7. K-C. Ong, A.W-K. Ng, L.S-U. Lee, G. Kaw, S-K. Kwek, M.K S. Leow, A. Earnest. Pulmonary function and exercise capacity in survivors of severe acute respiratory syndrome. Eur Respir J Sep 2004;24(3):436–442; doi: 10.1183/09031936.04.00007104.

8. Farsalinos K, Barbouni A, Niaura R. Systematic review of the prevalence of current smoking among hospitalized COVID-19 patients in China: could nicotine be a therapeutic option? Intern Emerg Med. 2020 May 9. doi: 10.1007/s11739-020-02355-7.

9. Farsalinos, K., Barbouni, A., Poulas, K., Polosa, R., Caponnetto, P., & Niaura, R. (2020). Current smoking, former smoking, and adverse outcome among hospitalized COVID-19 patients: a systematic review and meta-analysis. Therapeutic Advances in Chronic Disease. https://doi.org/10.1177/2040622320935765.

10. CDC COVID-19 Response Team. Preliminary estimates of the prevalence of selected underlying health conditions among patients with coronavirus disease 2019 — United States, February 12–March 28, 2020. MMWR Morb. Mortal. Wkly. Rep. 69, 382–386 (2020). https://doi.org/10.15585/mmwr.mm6913e2.

11. Hippisley-Cox J, Young D, Coupland C, et al Risk of severe COVID-19 disease with ACE inhibitors and angiotensin receptor blockers: cohort study including 8.3 million people Heart Published Online First: 31 July 2020. doi: 10.1136/heartjnl-2020-317393

12. Rossato M, Russo L, Mazzocut S, Di Vincenzo A, Fioretto P, Vettor R. Current Smoking is Not Associated with COVID-19. Eur Respir J. 2020 Jun; 55(6): 2010. doi: 10.1183/13993003.01290-2020.

13. Farsalinos K, Barbouni A, Niaura R. Smoking, vaping and hospitalization for COVID-19. Qeios ID: Z69O8A.11. 2020b. doi. 10.32388/Z69O8A.11.

14. Farsalinos K, Niaura R, Le Houezec J, Barbouni A, Tsatsakis A, Kouretas D, Vantarakis A, Poulas K. Editorial: Nicotine and SARS-CoV-2: COVID-19 may be a disease of the nicotinic cholinergic system. Toxicol Rep. 2020 Apr 30. doi: 10.1016/j.toxrep.2020.04.012.

15. Tracey KJ. The inflammatory reflex. Nature 2002; 420:853–9. doi: 10.1038/nature01321.

16. Wang H, Yu M, Ochani M, Amella CA, Tanovic M, Susarla S, Li JH, Wang H, Yang H, Ulloa L, Al-Abed Y, Czura CJ, Tracey KJ. Nicotinic acetylcholine receptor alpha7 subunit is an essential regulator of inflammation. Nature. 2003 Jan 23;421(6921):384–8.

17. Fujii T, Mashimo M, Moriwaki Y, Misawa H, Ono S, Horiguchi K, Kawashima K. Expression and Function of the Cholinergic System in Immune Cells. Front Immunol. 2017 Sep 6;8:1085. doi: 10.3389/fimmu.2017.01085.

18. Farsalinos K, Eliopoulos E, Leonidas D, Papadopoulos G, Tzartos S, Poulas, K. Molecular Nicotinic Cholinergic System and COVID-19: In Silico Identification of an Interaction between SARS-CoV-2 and Nicotinic Receptors with Potential Therapeutic Targeting Implications. Int. J. Mol. Sci. 2020:21(16);5807. doi: 10.3390/ijms21165807.

19. Oliveira, A., Ibarra, A. A., Bermudez, I., Casalino, L., Gaieb, Z., Shoemark, D. K., Gallagher, T., Sessions, R. B., Amaro, R. E., & Mulholland, A. J. (2020). Simulations support the interaction of the SARS-CoV-2 spike protein with nicotinic acetylcholine receptors and suggest subtype specificity. bioRxiv 2020.07.16.206680. doi: 10.1101/2020.07.16.206680.

20. Yuan M, Wu NC, Zhu X, Lee CD, So RTY, Lv H, Mok CKP, Wilson IA. A highly conserved cryptic epitope in the receptor binding domains of SARS-CoV-2 and SARS-CoV. Science. 2020 May 8;368(6491):630–633. doi: 10.1126/science.abb7269.

21. Huo J, Zhao Y et al, Neutralization of SARS-CoV-2 by destruction of the prefusion Spike Cell Host & Microbe doi: 10.1016/j.chom.2020.06.010

22. X. Tian, C. Li, A. Huang, S. Xia, S. Lu, Z. Shi, L. Lu, S. Jiang, Z. Yang, Y. Wu, T. Ying, Potent binding of 2019 novel coronavirus spike protein by a SARS coronavirus-specific human monoclonal antibody Emerging Microbes & Infections, 9:1, 382–385, DOI: 10.1080/22221751.2020.1729069

23. H Liu, NC. Wu, M Yuan, S Bangaru, JL. Torres, TG Caniels, J van Schooten, X Zhu, Chang-Chun D. Lee, Philip J.M. Brouwer, Marit J. van Gils, Rogier W. Sanders, Andrew B. Ward, Ian A. Wilson. Cross-neutralization of a SARS-CoV-2 antibody to a functionally conserved site is mediated by avidity bioRxiv 2020.08.02.233536; doi: https://doi.org/10.1101/2020.08.02.233536

24. Tsetlin VI, Hucho F. Snake and snail toxins acting on nicotinic acetylcholine receptors: fundamental aspects and medical applications. FEBS Lett. 2004 Jan 16;557(1-3):9–13.

25. Kalamida D, Poulas K, Avramopoulou V, Fostieri E, Lagoumintzis G, Lazaridis K, Sideri A, Zouridakis M, Tzartos SJ. Muscle and neuronal nicotinic acetylcholine receptors. Structure, function and pathogenicity. FEBS J. 2007 Aug;274(15):3799–845.

26. Basic Local Alignment Search Tool. Available online: http://www.ncbi.nlm.nih.gov/BLAST/

27. UniProt. UniProtKB - P0DTC2 (SPIKE_SARS2).

28. Sievers F, Wilm A, Dineen D, Gibson TJ, Karplus K, Li W, Lopez R, McWilliam H, Remmert M, Soding J.; et al. Fast, scalable generation of high-quality protein multiple sequence alignments using Clustal Omega. Mol. Syst. Biol. 2011, 7, 539.

29. Bender, B. J., Cisneros, A., 3rd, Duran, A. M., Finn, J. A., Fu, D., Lokits, A. D., Mueller, B. K., Sangha, A. K., Sauer, M. F., Sevy, A. M., Sliwoski, G., Sheehan, J. H., DiMaio, F., Meiler, J., & Moretti, R. (2016). Protocols for Molecular Modeling with Rosetta3 and RosettaScripts. Biochemistry, 55(34), 4748–4763. https://doi.org/10.1021/acs.biochem.6b00444

30. Gydo C P van Zundert, Alexandre M J J Bonvin. Modeling Protein-Protein Complexes Using the HADDOCK Webserver “Modeling Protein Complexes with HADDOCK”. Methods in molecular biology (Clifton, N.J.) 01/2014; 1137:163–79. DOI: 10.1007/978-1-4939-0366-5_12.

31. Li C. Xue, João Pglm Rodrigues, Panagiotis L. Kastritis, Alexandre Mjj Bonvin, Anna Vangone, PRODIGY: a web server for predicting the binding affinity of protein–protein complexes, Bioinformatics, Volume 32, Issue 23, 1 December 2016, Pages 3676–3678, https://doi.org/10.1093/bioinformatics/btw514

32. Goddard TD, Huang CC, Ferrin TE. Software extensions to UCSF chimera for interactive visualization of large molecular assemblies. Structure. 2005;13(3):473–482. doi: 10.1016/j.str.2005.01.006.

33. Chasapis CT, Kandias NG, Episkopou V, Bentrop D, Spyroulias GA. NMR-based insights into the conformational and interaction properties of Arkadia RING-H2E3 Ub ligase. Proteins, 80(5), 2012, 1484–1489.

34. Vaduganathan M, Vardeny O, Michel T, McMurray JJV, Pfeffer MA, Solomon SD. Renin-Angiotensin-Aldosterone System Inhibitors in Patients with Covid-19 N Engl J Med. 2020;382(17):1653–1659. doi: 10.1056/NEJMsr2005760

35. Alcami, A., and Koszinowski, U. H. (2000) Viral mechanisms of immune evasion. Immunol. Today 21, 447–455. N EnglJ Med. 2020. doi: 10.1056/NEJMsr2005760

36. Duffy, S., Shackelton, L. A., and Holmes, E. C. (2008) Rates of evolutionary change in viruses: patterns and determinants. Nat. Rev. Genet. 9, 267–276.

37. Wang, L., Parr, R. L., King, D. J., and Collisson, E.W. (1995) A highly conserved epitope on the spike protein of infectious bronchitis virus. Arch. Virol. 140, 2201–2213.

38. Sahini, L., Tempczyk-Russell, A., and Agarwal, R. (2010) Large-scale sequence analysis of hemagglutinin of influenza A virus identifies conserved regions suitable for targeting an anti-viral response. PLoS One 5, e9268.

39. Ashkenazi, A., Faingold, O., Kaushansky, N., Ben-Nun, A., and Shai, Y. (2013) A highly conserved sequence associated with the HIV gp41loop region is an immunomodulator of antigen-specific T cells in mice. Blood 121, 2244–2252.

40. Burton, D. R., Poignard, P., Stanfield, R. L., and Wilson, I. A. (2012) Broadly neutralizing antibodies present new prospects to counter highly antigenically diverse viruses. Science 337, 183–186.

